# Chorus Synchrony as a Cue: Influence of Chorus Howl Synchrony from Unfamiliar Packs on Wolves’ Behaviour

**DOI:** 10.64898/2026.08.20.745898

**Authors:** Vanessa Kern, Svenja Capitain, Gwendolyn Wirobski, Pablo Arias-Sarah, Martha Newson, Erica van de Waal, Friederike Range

## Abstract

Chorus howls play a central role in wolves’ (*Canis lupus*) social communication, shaping both intra- and inter-pack interactions. While synchrony has been shown to influence perceived threat from outgroups in birds and humans, whether the synchrony of an outside pack’s chorus howl plays a role in wolf pack behaviour remains unclear. This study investigated behavioural responses of 19 zoo-housed wolf packs to playbacks of synchronous and asynchronous chorus howls from unfamiliar packs, in comparison to a dove control call. We hypothesized that packs would show stronger intra-pack cohesion and greater spatial withdrawal in response to more synchronous, and thus cohesive – and potentially more formidable – sounding vocalisations. Alternatively, asynchronous howls might elicit stronger responses due to their ambiguity and the “Beau Geste” effect (where asynchronous signals make it harder to infer the number of callers). Synchronous howls resulted in more distance from the sound source and less activity compared with asynchronous howls. This could be interpreted as a more cautious response towards a perceived threat. However, since there was no effect on pack-directed behaviour and no difference between the synchronous and control sound, lower interest or motivation to engage with the stimulus cannot be excluded. In contrast, wolves spent more time near the sound-source after hearing asynchronous howls, approached it more, and oriented towards it for longer compared to synchronous playbacks, a pattern consistent with threat-investigation. In how far these reactions result from a more difficult localisation of synchronous howls or more ambiguity in pack size in the asynchronous playbacks, as well as how this translates to wild wolf behaviour, remains to be investigated.

---

Group coordination plays a critical role in the survival and success of social animals (Couzin & Krause, 2003; Sumpter, 2006). Across species, behaviours such as collective movement, ritualised displays, and vocalisations facilitate cooperation, signal social cohesion, and influence social interactions both within and between groups (Couzin & Krause, 2003; Sumpter, 2006). Especially coordinated acoustic signals have evolved as a central means of communication in many group-living species, including canines and primates (Harrington & Asa, 2003; Snowdon, 2017), mediating intra-group bonding and, at the same time, influencing inter-group dynamics (Couzin & Krause, 2003; Sumpter, 2006). Long-distance acoustic signals, in particular, often serve dual functions. They may facilitate group reunification, such as in response to separation or threat, and simultaneously function as displays of strength or territoriality toward outgroups (Byrne, 1982; Harrington & Asa, 2003). The structure and synchrony of these signals can vary, providing differential information to receivers (Kinzey & Robinson, 1983).

In several species, there is behavioural evidence that higher degrees of coordination, i.e., more synchronous acoustic signals, may be perceived as more threatening or impressive than asynchronous ones, especially by outgroup members (Bortolini et al., 2025; Hall & Magrath, 2007). For example, in bird species, studies on breeding pairs have shown that synchronised vocal duets may be perceived as more threatening as they were more effective for territorial defence than asynchronous displays (Hall & Magrath, 2007). Similarly, in humans, synchronized movement and sound can increase perceived group strength and cohesion. The sound of military groups marching in synchrony is judged to be more formidable (i.e. perceived capacity to succeed in physical confrontations or pose a credible threat to others) than marching asynchronously, a perception mediated both by the group appearing as a unified whole (entitativity) and by inferences of strong internal bonds (Fessler & Holbrook, 2016). Comparable effects have also been observed in non-institutional settings, where coordinated football chants have been associated with heightened in-group cooperation, increased hostility toward outgroups and greater perceived group formidability (Bortolini et al., 2025, Newson et al., in review). Furthermore, more harmonious signals seem to make it more difficult for human observers to assess what direction the auditory signal came from (Bregman, 1990; Harrington & Asa, 2003).

In contrast, asynchronous or highly modulated group signals have been proposed to increase perceptual ambiguity regarding the number of callers, which could exaggerate the apparent size and strength of the signalling group (Bee & Micheyl, 2008; Brewster et al., 2017; Filibeck et al., 1982; Harrington, 1989; Harrington & Asa, 2003; Newson et al., in review; Wiley & Richards, 1978). Coined as the *Beau Geste effect,* the deception strategy resulting from this form of temporal asynchrony has originally been described in birds by Krebs (1977). He observed that territorial birds that use large and varied song repertoires create the impression of multiple competitors, thereby exaggerating the apparent density of rivals (Krebs, 1977). In line with this, a recent paper comparing UK participants’ reaction to culturally non-salient synchronous and asynchronous stimuli (Brazilian football chants) found that asynchronous chants were perceived as more threatening than synchronous ones (Newson et al., in review). Further support comes from analyses of wolf and coyote howling showing that, when calls lack stable temporal alignment, differentiating between individual acoustic tracks becomes more difficult (Brewster et al., 2017; Filibeck et al., 1982; Harrington, 1989; Harrington & Asa, 2003). As a result, even experienced human listeners and automated acoustic analyses fail to accurately estimate the number of callers (Brewster et al., 2017; Filibeck et al., 1982; Harrington, 1989; Harrington & Asa, 2003). However, in how far this affects intraspecific behaviour has not yet been tested in canids.

Wolves present an intriguing case to test these hypotheses. Long-distance vocalisations play a central role in their social ecology, with chorus howls, which involve two or more wolves, serving multiple functions at both the intra- and inter-pack level. Within packs, chorus howling contributes to group cohesion, social bonding and reunification, particularly following partial separation or arousal-inducing events (Harrington & Mech, 1979; Nowak et al., 2007; Theberge & Theberge, 2022). At an inter-pack level, chorus howls function as long-distance territorial signals that facilitate spacing between neighbouring packs while reducing the need for direct physical encounters (Harrington & Mech, 1979; Nowak et al., 2007; Theberge & Theberge, 2022). Decades of evidence have shown that wolves adjust to incoming howls from outgroups depending on their own pack’s social context. For example, larger packs are more likely to remain in place and respond vocally to howls from unfamiliar conspecifics, likely due to their higher competitive ability and reduced risk association with inter-pack encounters (Cubaynes et al., 2014; Harrington & Asa, 2003; Harrington & Mech, 1979). Additionally, the presence and age of more dominant individuals, especially older animals and males, have been associated with an increased likelihood of vocal responses and reduced retreat behaviour (Cassidy et al., 2015; Cubaynes et al., 2014; Harrington & Asa, 2003; Harrington & Mech, 1979; Mech & Boitani, 2003). Despite this evidence that packs adjust their responses to chorus howls based on their own social context, it remains unclear whether wolves attend to the acoustic features of incoming wolf howls.

This is particularly interesting given that wolf howls show flexibility in acoustic structure, timing and coordination. Schassburger (1987) described the existence of two distinct types of chorus howls, potentially representing opposite ends of an acoustic spectrum: harmonious howls, where the group howls show little modulation between individuals, and discordant howls, characterised by less coordinated and more irregular patterns (Harrington & Asa, 2003; Palacios et al., 2007). However, it remains unclear to what extent variation in synchrony modulates how information about the signalling group is perceived by receivers, and how this is influenced by the characteristics of the receiving group.

Thus, the present study investigated whether zoo-housed wolves exhibit different behavioural responses to synchronous compared to asynchronous chorus howl playbacks from an unfamiliar pack. To control for potential effects of sound transmission through the loudspeaker, including changes in acoustic quality and perceived naturalness of the sound, and to have a baseline of the wolves’ behaviour, a neutral heterospecific control stimulus (dove call) was included in the within-subject design. Our first hypothesis postulates that synchronized howls signal greater intra-pack cohesion and therefore higher group formidability, as was previously shown in humans and birds (Bortolini et al., 2025; Fessler & Holbrook, 2016; Hall & Magrath, 2007). Accordingly, we predicted that wolves would respond with increased avoidance of the sound-source and stronger intra-pack cohesion, such as increased physical proximity with conspecifics, affiliative behaviour and/or orientation towards each other. These responses might be enforced by reduced localizability of more harmonious howls (Harrington & Asa, 2003), potentially linked to less gazing towards the sound-source compared to the asynchronous playback. Alternatively, we hypothesised that the *Beau Geste* effect may take effect in inter-pack communication, in which asynchronous and discordant signals may make it harder to infer the number of vocalising individuals (Brewster et al., 2017; Filibeck et al., 1982; Harrington & Asa, 2003). In this case, wolves would be expected to exhibit stronger avoidance of the sound-source and greater intra-pack cohesion in response to asynchronous rather than synchronous howls. Furthermore, we expected, independently of the chorus howl condition, reduced avoidance of the sound-source with increased pack size, higher mean group age, and higher male-female ratio in the group. Finally, if wolves respond specifically to the howl stimuli rather than to the loudspeaker playback itself, stronger and clearer behavioural responses were expected in both howl conditions compared to the control call.

## MATERIAL & METHODS

### Ethical Statement

All procedures complied with institutional and national animal-ethics regulations. Ethical approval was granted from the Ethics and Animal Welfare Committee of the University of Veterinary Medicine, Vienna in accordance with the University’s guidelines for Good Scientific Practice (ETK-078/07/2024 and ETK-063/05/2025). The study was carried out under a valid Swiss animal-testing licence (SZ-38018/2025) in accordance with the Swiss Animal Welfare Act (TSchG, SR 455, Kap.3, Art. 22). Additionally, all site-specific details were discussed and approved with the zoo staff.

### Study Animals and Location

Nineteen captive grey wolf packs from fifteen zoos across Austria and Switzerland took part in the study, distributed across two separate field seasons in 2024 and 2025. Each pack consisted of two to seven animals (80 individuals in total). The animals were all tested as a group in their outdoor home enclosures in each zoo.

### Stimuli

The synchronous and asynchronous howling tracks used in this study were selected from material previously recorded by Vlasitz (2022) at the Core Facility Wolf Science Center in Ernstbrunn, Austria (see Mazzini et al., 2013; Watson et al., 2018 for similar data collection). Six dyadic howls were selected: three naturally synchronous and three naturally asynchronous. Both howl types were obtained from the same three packs (each consisting of two individuals), with each pack contributing one synchronous and one asynchronous howl. These three howl pairs were chosen because they had the highest mutual information distance (i.e., howl-to-howl similarity among the two howling individuals) between synchronous and asynchronous tracks in each pair, while matching in general acoustic properties such as average frequency, range, and coefficient of variation.

To quantify howl-to-howl similarity we used Mutual Information. Specifically, we computed the degree of zero-lag vocal synchrony between interacting individuals by computing the Mutual Information (MI) between time-aligned pitch contours. Here, MI was used to represent the amount of shared information – or dynamic structural similarity – between the two co-occurring and temporally overlapping vocalization time series, capturing both linear and non-linear coordinated frequency modulations (Shahsavari Baboukani et al., 2020). We estimated MI using the Kraskov-Stögbauer-Grassberger (KSG) nearest-neighbour algorithm (Kraskov et al., 2004). To compute MI, we transformed the pitch contours to cents and used the overlapping pitch contours to compute MI using a neighbourhood size of k = 4 and a base-2 logarithm. Final MI values are in bits, and higher values indicate stronger synchronisation between the two pitch contours. As expected, across the whole dataset, this measure of Mutual Information significantly correlated with maximum and mean cross correlation measures of synchronisation (max cross-correlation correlation vs MI: *P* < 0.001, r = 0.410; mean cross-correlation correlation vs MI: *P* < 0.001, r = 0.400; Fig. S1). Synchronous howls had significantly higher Mutual Information than asynchronous howls.

Since the recordings only contained howling bouts (ranging from 7-13s) rather than full-length howling sessions (those are generally interrupted by background noises and breaks), the playback stimuli were generated by concatenating the same bout to a length of approximately 17 seconds. Not only did this standardise the duration of the stimuli across conditions, but it also represented a more natural howling session length (Harrington, 1989; Harrington & Asa, 2003). This procedure was applied identically to synchronous and asynchronous stimuli to avoid introducing systematic differences in stimulus construction. Potential pitfalls of this playback generation process are addressed in the discussion.

To control for potential effects of the loudspeaker playback itself, a control condition was included, using a song call from a single Eurasian collared dove (*Streptopelia decaocto*) matched in loudness and duration to the wolf stimuli (source: Xeno-canto.org, Francesco Sottile). The locations and habitats of the tested wolves overlapped with the distribution of the Eurasian collared dove, a widespread species in Europe (Romagosa & Mlodinow, 2020). Hence, it is very likely that the wolves were familiar with this type of call from the surrounding woods, making it an ideal control sound.

All playback stimuli were broadcast using a LD Systems Roadboy 65 loudspeaker at a maximum sound pressure level of 110 dB SPL (measured at 1 m).

### Procedure

Data collection was conducted between September and December to avoid the mating season (typically late December to April) during which intra-pack behaviour differs from typical year-round patterns (Harrington & Mech, 1979). We further excluded the early months following pup births (April to August), when howling response rates are particularly low (McIntyre et al., 2017; Nowak et al., 2007). Since previous studies have shown that wolves are most likely to howl during the evening hours, particularly in response to elicited howls in captive settings (Nowak et al., 2007), our tests were scheduled between 17.00 and 18.00 during the warmest months and 15.00 and 16.00 during cooler periods to account for changes in daylight hours. We coordinated with the zookeepers to schedule the test sessions on days that were similar in terms of routine conditions, like feeding times or visitor programs.

Each of the three playback conditions (synchronous howl, asynchronous howl (both originating from the same pack), and control dove) were presented once per pack in a within-subject design. A minimum interval of three weeks (average interval: 22.3 days; range: 21 – 42 days) was maintained between sessions to minimize carry-over effects. The order of the playback conditions was randomized and counterbalanced across all groups.

During the first season of data collection (year 2024), an initial observation session was conducted a day prior to the first playback test days, wherein the animals were observed and recorded for a period of approximately one hour. This session served as a baseline for comparison for the termination criteria in later sessions, as well as to determine optimal placement for the loudspeaker and cameras. For logistical reasons, this procedure was carried out one hour before the first playback session during the second season (year 2025) of data collection. Since we did not directly interact with the animals during this period and did not walk off the visitor path, it is unlikely that this change affected animal behaviour during the playback sessions.

On playback test days, four cameras (Sony Handycam camcorders) were set up around the enclosures to maximize coverage of the area: three at fixed locations around the enclosure, including one near the loudspeaker, and one handheld by the experimenter. The loudspeaker was placed 1-2 metres from the enclosure fence, behind vegetation, outside the visitor area, i.e. in locations with minimal or no human presence, to minimize the association of the sound with humans. The camera and loudspeaker positions remained consistent for all subsequent sessions for the respective pack. The cameras and loudspeaker were switched on and the experimenter moved to their viewing position on the visitor path to ensure that their movement was clearly separated from the playback itself. Each playback track contained an initial five-minute silence, so the howl was broadcasted five minutes after the loudspeaker was activated. Once the playback started, the session began and the animals were filmed for 15 minutes. According to the protocol, any prolonged agitation or withdrawal would have required the termination of the current and future sessions. However, no such behaviour was observed.

### Measured Behaviours

The videos were coded using BORIS software (v. 9.3.2, Friard & Gamba, 2016). The ethogram included behaviours directed at the sound-source, behaviours directed towards conspecifics, vocalisations, self-directed behaviours, and general activity (Table 1). Because individuals within each group could not be reliably identified across the three playback days, all animals in the videos were coded individually, but their behaviour was later averaged across all individuals per pack and playback session. This provided us with a measure of the average behaviour (for each behavioural variable respectively) per pack for each playback session, which could be used as analysis unit with reliable control for the repeated design (Group as random factor, see below). All videos were coded blind to condition and inter-observer reliability was assessed on 20.4 % of the dataset (11 videos) by a second coder who was blind to the condition, achieving an interrater agreement of 0.89 (ICC 0.70-1.00).

**Table 1.**
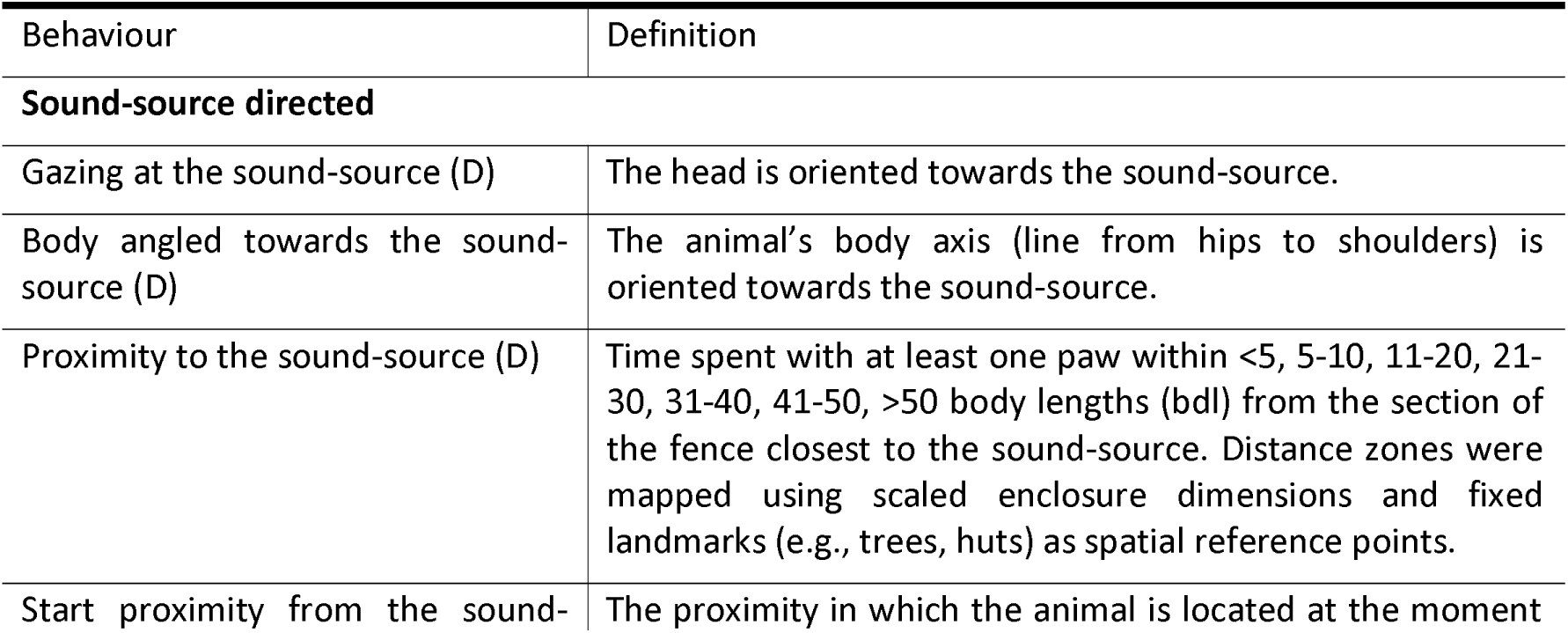
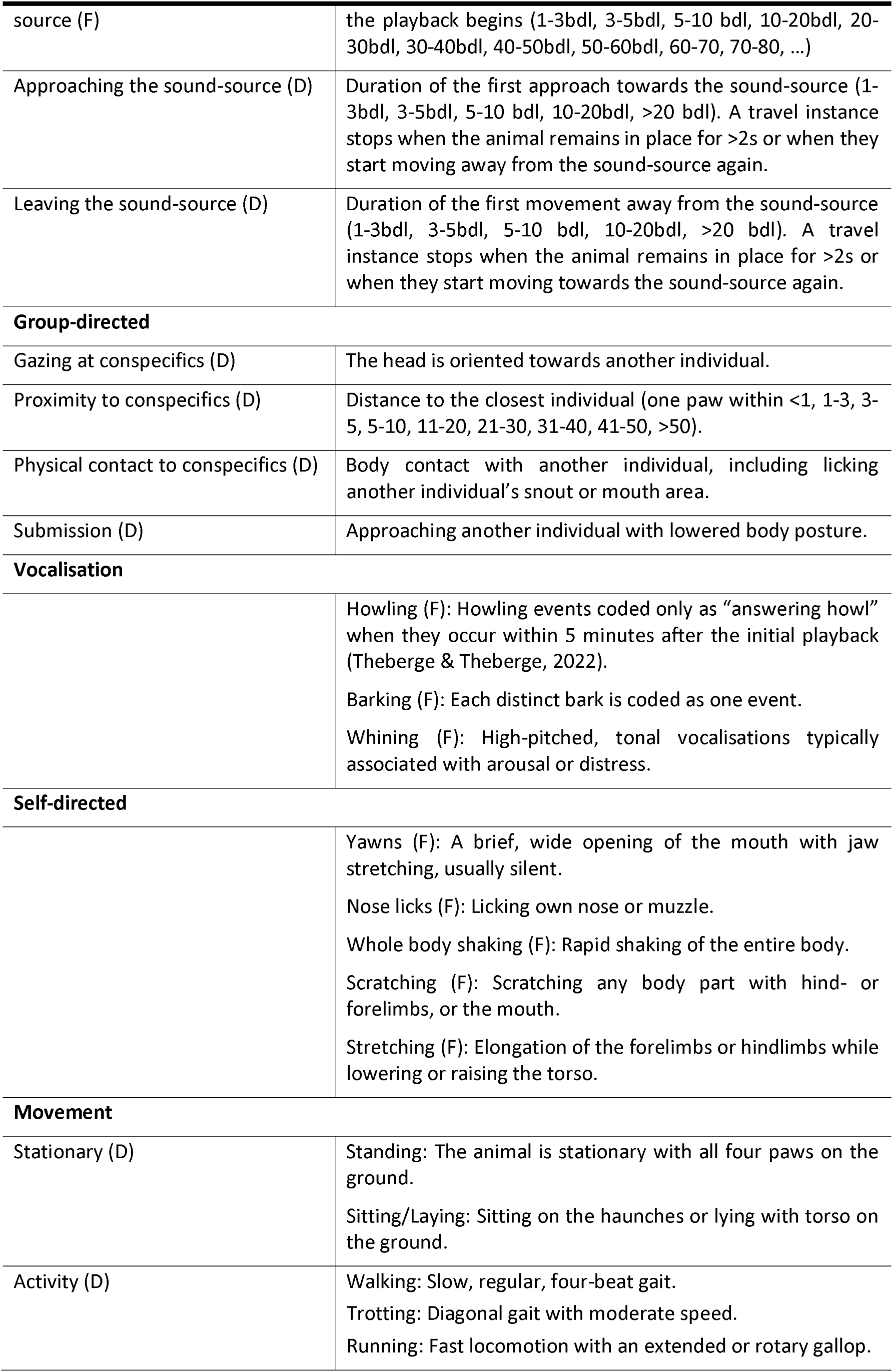
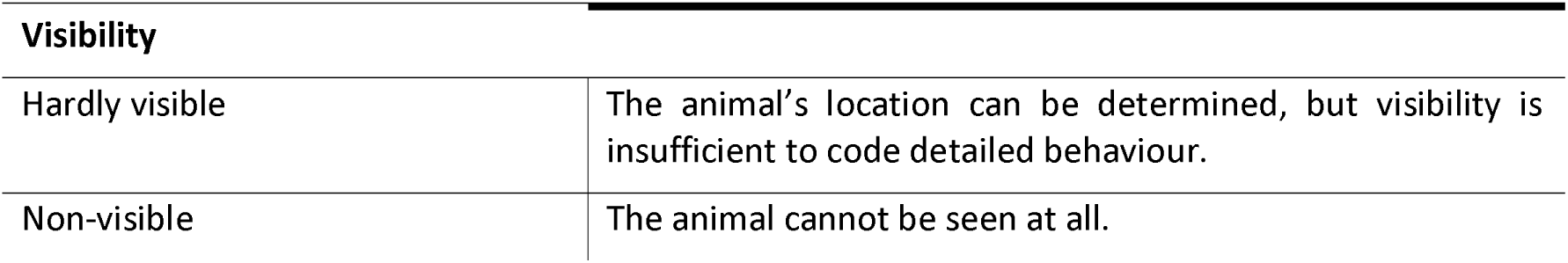
Ethogram of all measured behaviours, including definitions and coding criteria. Behaviours were coded as durations (D) or frequencies (F).

### Statistical Analysis

Statistical analyses were conducted in R (R Studio v4.4.2; R Core Team, 2024). To ensure data reliability, observations in which individuals were not visible for more than 70% of the test duration were excluded from the analysis (n = 46 out of 171 individual observations, n = 8 out of 51 group averages). Variables coded as durations were analysed as proportions of the time animals were visible. Infrequent behaviours were changed into binary values (approach sound, leave sound, self-directed behaviours) for each observation and behaviours that occurred in less than 10% of trials were excluded from the analysis (physical contact, licking, submission, vocalisations). For the analyses, each 15-min observation was divided into three 5-min intervals to assess behavioural responses over time.

For behaviours expressed as proportions of the total test duration, we fitted generalized linear mixed models using the glmmTMB package in R (Brooks et al., 2017). Since beta models cannot accommodate exact values of 0 or 1, proportional response variables were transformed prior to analyses. Numeric predictors were z-transformed before model fitting to improve model fit and interpretability. Playback condition was included as the main predictor, while time period (0-5 min, 5-10 min, 10-15 min post-playback), pack-average starting proximity to the sound- source at the onset of the playback, playback day, pack size and sex ratio (number of females divided by the total number of females and males) were included as control predictors. Age was initially considered as an additional control predictor but was excluded because exact age data were not available for all individuals. Group ID was included as a random effect to account for the repeated-measures design.

Distance variables were originally coded as the duration spent within predefined distance categories (e.g., <5 body lengths (bdls), 5-10 bdls, … >50 bdls). To obtain a single continuous distance measure for each observation, we calculated a time-weighed mean distance. For each distance category, the duration spent in that category was multiplied by the midpoint of the respective distance range (e.g., 7 for the 5-10 bdls, 15 for the 11-20 bdls category). These weighted values were then summed across distance categories and divided by the total duration for which distance was recorded. Since proximity to conspecifics was right-skewed, it was log-transformed prior to analyses to improve normality. The shortest distance to the sound- source over the entire observation period was also based on the midpoint of the lowest distance category. Distance variables were then analysed using linear mixed models fitted with the lmer function (lme4 package in R; Bates et al., 2015) following the same structure as described above. When full-null model comparisons were significant, full models were refitted by means of the Satterthwaite approximation using restricted maximum likelihood estimation (REML; Luke, 2017).

Model assumptions, collinearity, and model stability were checked for each model, and each full model was compared with its corresponding null model lacking the main test predictors to test the overall contribution of the fixed effects. To reduce the likelihood of Type I errors, model interpretation only proceeded if this comparison was significant. Drop-one likelihood-ratio tests using the drop1 function (R Core Team, 2024) were used to identify significant predictors, and post hoc pairwise comparisons for significant categorical predictors were conducted using estimated marginal means with Tukey-adjusted p-values (emmeans package; Lenth & Piaskowski, 2017). Confidence intervals for fixed effects were obtained via parametric bootstrapping.

## RESULTS

### Sound-directed Behaviours

The mean distance of the group to the sound-source across the entire playback duration was significantly affected by playback condition (*F*_2, 102.957_ = 5.352, *P* = 0.006). At the group level, distance to the sound-source was significantly lower during the asynchronous condition than during the synchronous condition (β = -4.800, SE = 1.470, t_103_ = -3.259, *P* = 0.004; Fig. 1). The asynchronous and control conditions did not differ significantly (β = -2.660, SE = 1.450, *t*_104_ = - 1.839, *P* = 0.162), nor did the difference between the control and the synchronous condition (β = -2.140, SE = 1.460, *t*_104_ = -1.464, *P* = 0.313). The mean distance to the sound-source increased significantly with group size (*F*_1, 14.510_ = 6.074, *P* = 0.027).

**Figure 1.**
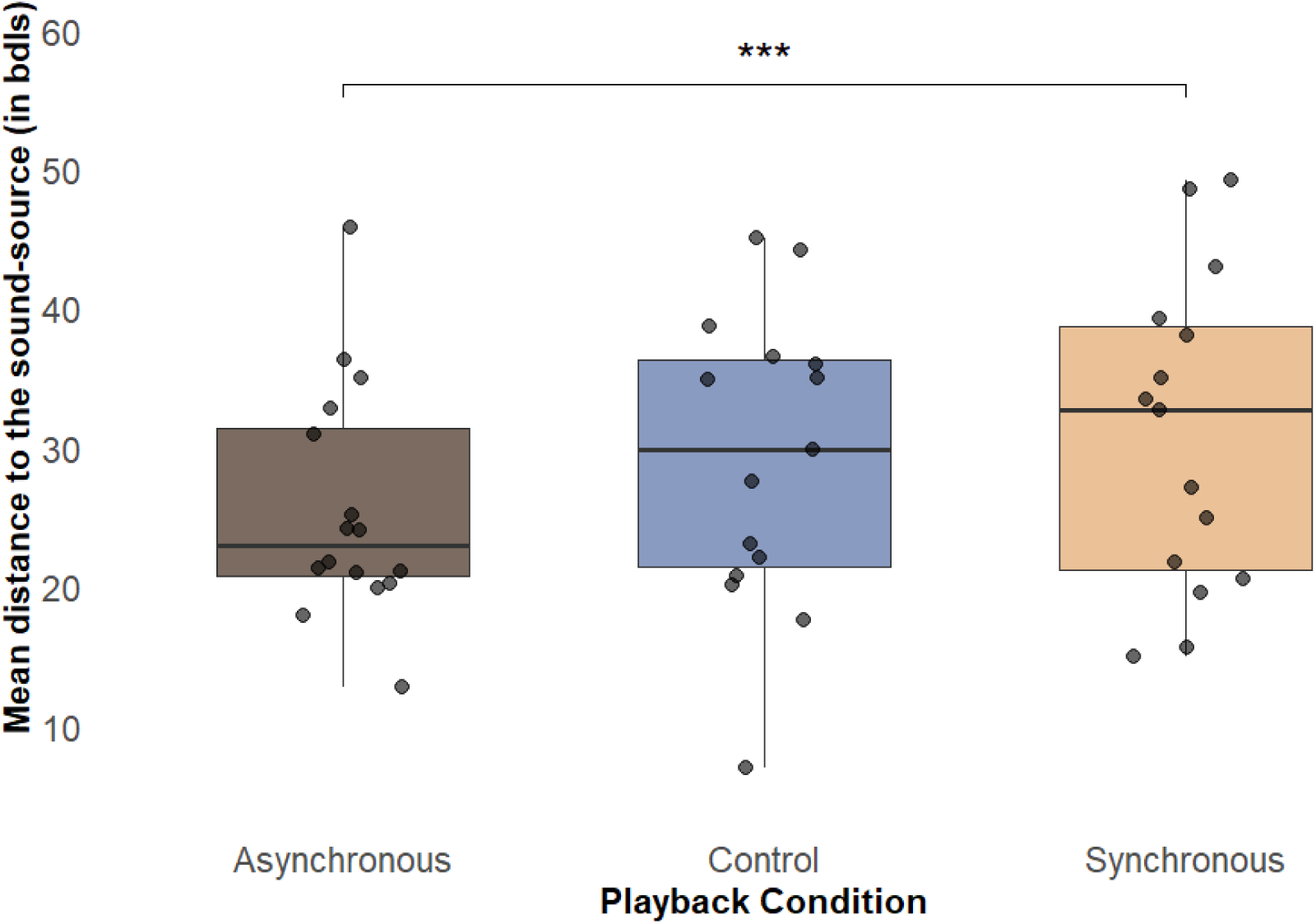
Mean distance (in body lengths) to the sound-source across playback conditions. Boxplots show wolves’ mean distance (averaged per group) to the sound-source in body lengths while visible during the 15-minute observation period after asynchronous chorus howls, dove control calls, and synchronous chorus howls. Boxes show the interquartile range, horizontal lines indicate medians, whiskers show 1.5 x IQR, and points represent individual group observations. Asterisks indicate significant differences between playback conditions with p=0.10 > ° > 0.05 > * > 0.01 > ** > 0.001 > ***.

Closest distance of the group to the sound-source was likewise significantly affected by playback condition (*F*_2, 104.830_ = 8.819, *P* < 0.001). Descriptively, groups approached the sound-source closest during the asynchronous condition (mean ± SD = 17.80 ± 10.60 bdls), followed by the control condition (mean ± SD = 21.70 ± 11.90 bdls), and approached with the largest distance during the synchronous condition (mean ± SD = 25.90 ± 12.40 bdls). At the group level, closest distance to the sound-source was significantly lower during the asynchronous than during the synchronous condition (β = -8.040, SE = 1.920, t_104_ = -4.191, *P* < 0.001). There was also a weak trend for closest distance to be lower during the control condition than during the synchronous condition (β = -4.200, SE = 1.900, t_105_ = -2.212, *P* = 0.074). In addition, the mean closest distance tended to increase with group size, suggesting that larger groups did not approach the sound-source as closely as smaller groups (*F*_1, 15.002_ = 4.311, *P* = 0.055). Approach behaviour provided additional evidence for condition-related differences in response to the sound-source. The likelihood of approaching the sound-source was significantly affected by playback condition (χ^2^_2_ = 14.524, *P* < 0.001), with groups being significantly more likely to approach the sound-source during the asynchronous condition than during the synchronous condition (β = 2.990, SE = 1.020, *z* = 2.921, *P* = 0.009; Fig. 2). The groups were significantly less likely to approach the sound-source during the synchronous condition than during the control condition (β = 2.700, SE = 0.975, *z* = 2.767, *P* = 0.016). The likelihood of approach decreased significantly with time after the playback (χ^2^_1_ = 4.440, *P* = 0.035) and increased with higher starting distance to the sound-source (χ^2^_1_ = 9.151, *P* = 0.002). In addition, there was a significant negative effect of the number of playback day (χ^2^_1_ = 4.971, *P* = 0.026), indicating a decrease in the likelihood of approach across sessions.

**Figure 2.**
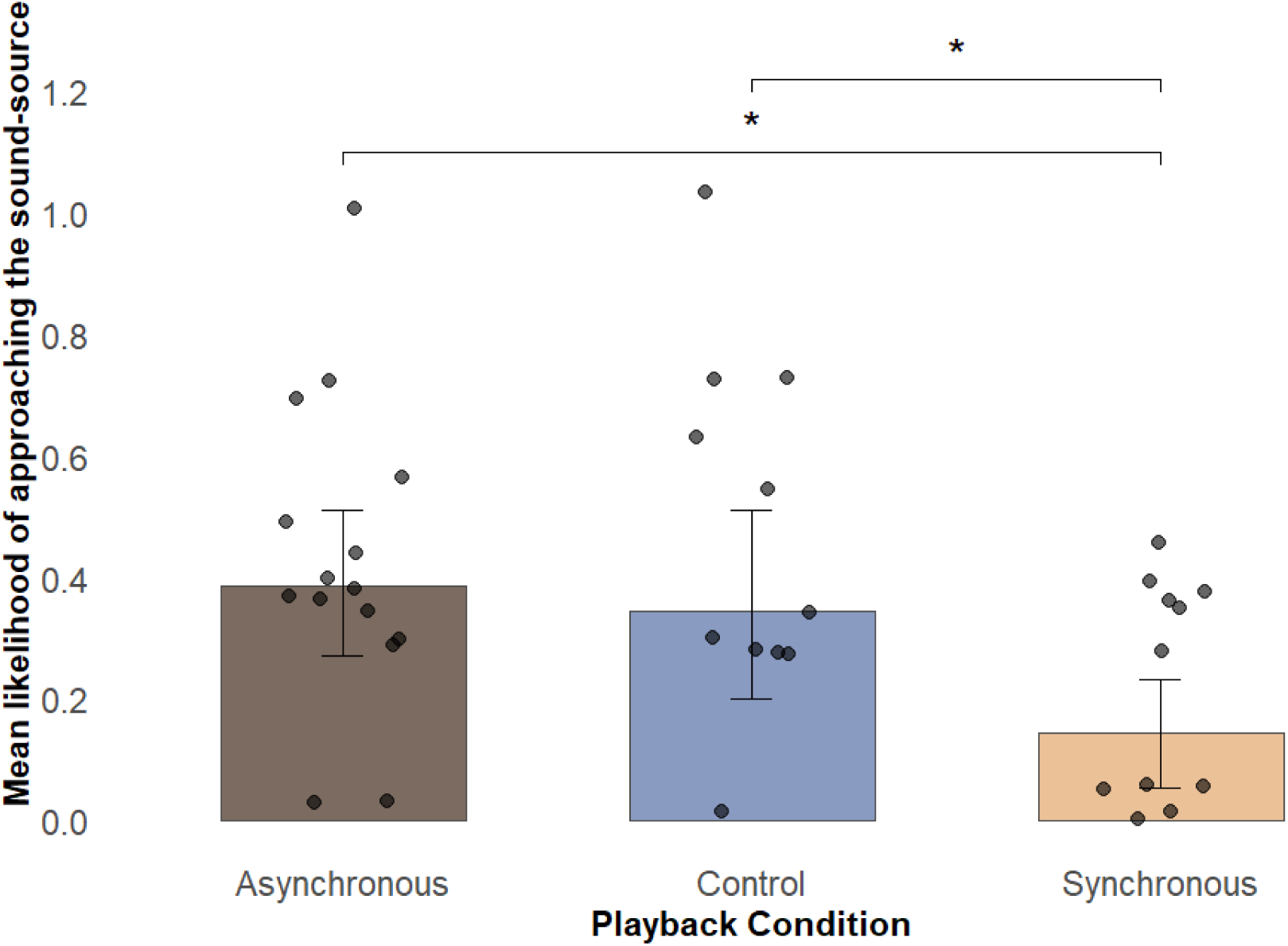
Mean likelihood of approach towards the sound-source across playback conditions. Approach was scored as a binary variable, with 0 indicating no approach and 1 indicating approach towards the sound-source during the 15-minute observation period. Bars show the mean approach score per pack following asynchronous chorus howls, dove control calls, and synchronous chorus howls. Error bars indicate 95% confidence intervals, and points represent individual group observations. Asterisks indicate significant differences between playback conditions with p=0.10 > ° > 0.05 > * > 0.01 > ** > 0.001 > ***.

For the likelihood to leave the sound-source, the full-null model comparison was not significant (χ^2^_4_ = 5.059, *P* = 0.281); therefore, individual predictors were not analysed further.

The overall effect of playback condition on gazing towards the sound-source showed a significant effect of condition (χ^2^_2_ = 6.015, *P* = 0.049). The group gazed towards the sound-source for longer during the asynchronous condition than during the synchronous condition, although the post-hoc effect only indicated a trend (β = 0.515, SE = 0.227, *z* = 2.267, *P* = 0.061). Gazing towards the sound-source also decreased significantly over time within each session (χ^2^_1_ = 5.733, *P* = 0.017) and increased with higher starting distance from the sound-source (χ^2^_1_ = 4.548, *P* = 0.033). Furthermore, groups with a higher proportion of males spent significantly more time gazing towards the sound-source than groups with higher female ratios (χ^2^_1_ = 5.210, *P* = 0.022).

Body orientation towards the sound-source showed only weak evidence for an effect of playback condition (χ^2^_2_ = 5.257, *P* = 0.072). Groups tended to spend more time oriented towards the sound-source during the asynchronous condition than during the control condition (β = 0.601, SE = 0.261, *z* = 2.300, *P* = 0.056). Sex ratio was also associated with body orientation towards the sound-source, with groups with a higher proportion of males spending significantly more time with their body angled towards the sound-source (χ^2^_1_ = 8.631, *P* = 0.003). Body orientation towards the sound-source also tended to decrease with higher starting distance from the sound-source (χ^2^_1_ = 3.314, *P* = 0.069). In addition, there was a weak negative effect of the number playback day (χ^2^_1_ = 3.052, *P* = 0.081), indicating a decrease in the likelihood of approach across sessions.

### General Behaviours

The level of activity was significantly affected by playback condition (χ^2^_2_ = 23.021, *P* < 0.001; Fig. 3). At the group level, there was more movement during the asynchronous condition than during the synchronous condition (β = 1.230, SE = 0.255, *z* = 4.832, *P* < 0.001), and during the control condition (β = 0.869, SE = 0.226, *z* = 3.849, *P* < 0.001).

**Figure 3.**
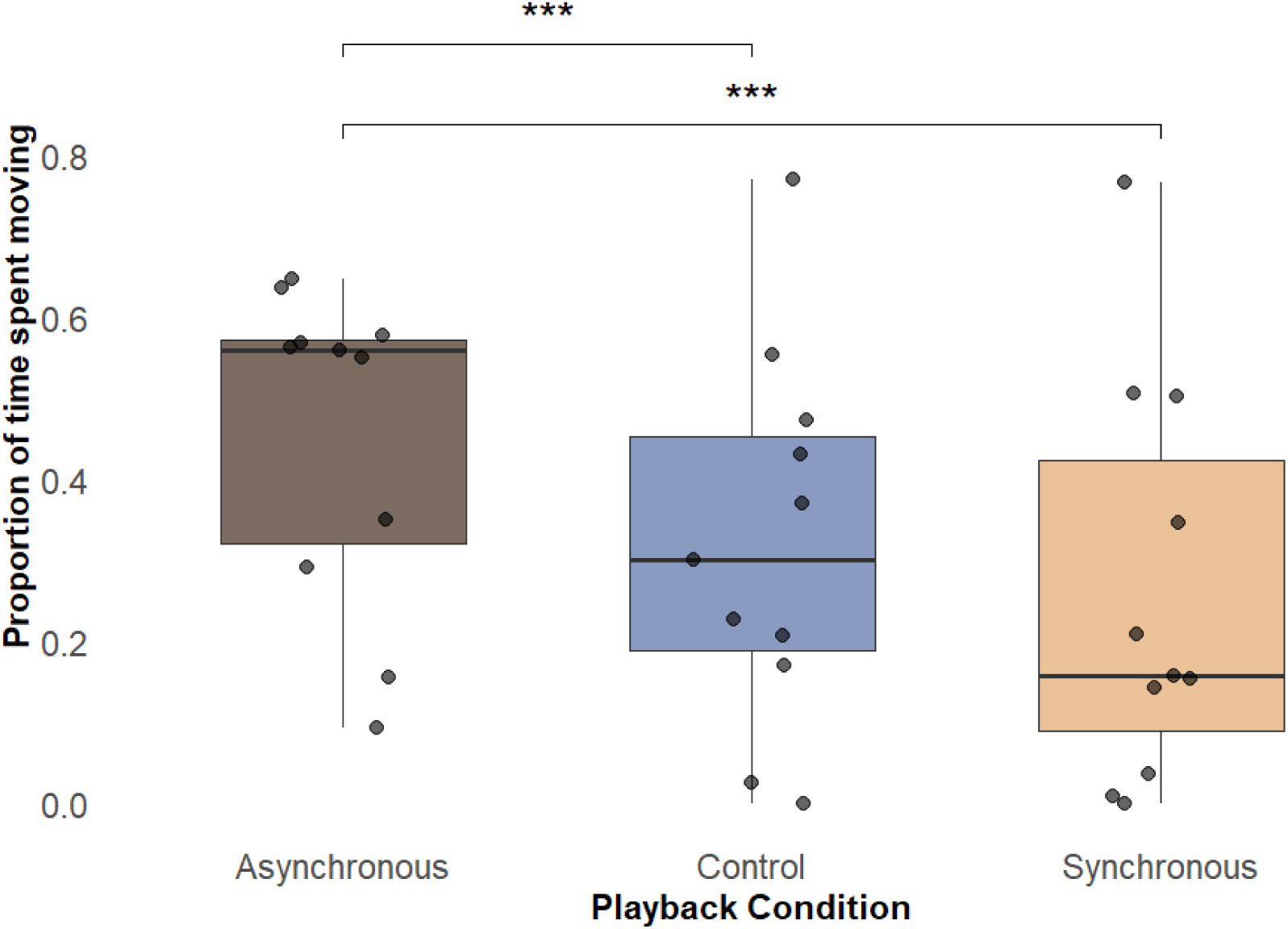
Time spent moving across conditions. The boxplots show the average proportion of time the packs spent moving during the 15-minute observation period following asynchronous chorus howls, dove control calls, and synchronous chorus howls. Boxes show the interquartile range, horizontal lines indicate medians, whiskers show 1.5 x IQR, and points represent individual group observations. Asterisks indicate significant differences between playback conditions with p=0.10 > ° > 0.05 > * > 0.01 > ** > 0.001 > ***.

Furthermore, sex ratio showed a positive association with the activity level, with groups containing a higher proportion of males spending less time moving (χ^2^_1_ =10.374, *P* = 0.001; Fig. 4).

**Figure 4.**
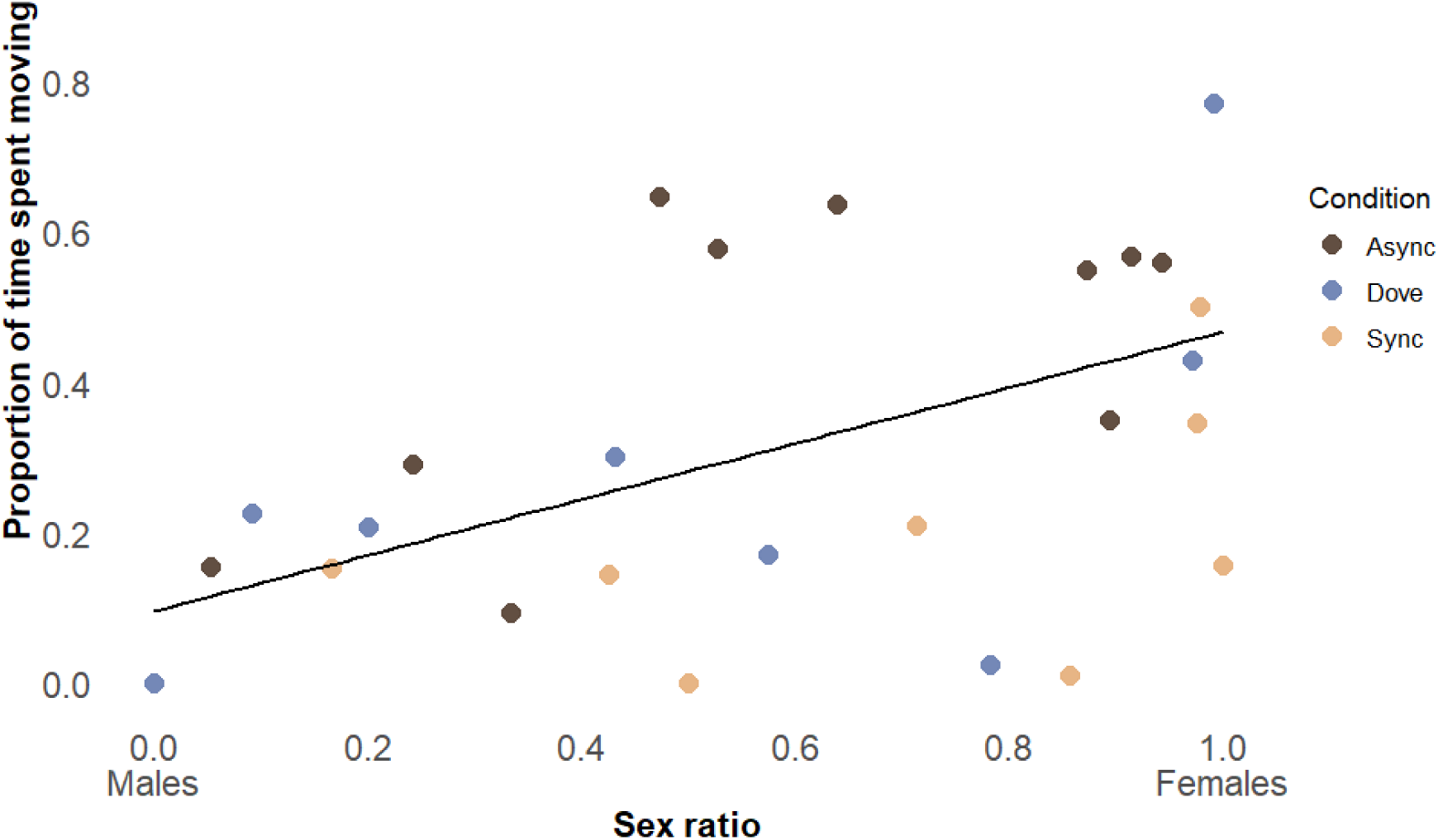
Relationship between group sex ratio and movement. The figure shows the proportion of time wolves spent moving during the 15-minute observation period in relation to the group sex ratio. Sex ratio was calculated as females / (females + males), with higher values indicating a higher proportion of females in the group. Points represent individual group observations, coloured by playback condition (asynchronized = brown, synchronized = yellow, control (dove) = blue); larger diamonds show the same observations at their exact values, while smaller jittered points are displayed to reduce overlap. Colours indicate playback condition. The black line shows the fitted regression line.

The full-null model comparison for self-directed behaviour showed a weak trend (χ^2^_4_ = 7.926, *P* = 0.094), but there was no evidence for significant differences between conditions in the post-hoc analysis. However, group size had a significant positive effect on self-directed behaviours, with larger groups being more likely to show self-directed behaviours (χ^2^_1_ = 4.264, *P* = 0.039). There was also a weak positive trend for start proximity, indicating that self-directed behaviours were more likely shown when individuals were farther away from the sound-source at the start of the playback (χ^2^_1_ = 3.193, *P* = 0.074).

### Group-directed Behaviours

For proximity to conspecifics, the full-null model comparison showed only weak evidence for improved model fit compared to the null model (χ^2^_4_ = 8.436, *P* = 0.077), and predictor-level effects should therefore be interpreted cautiously. However, within the full model, playback condition showed no evidence for an effect on proximity to conspecifics (*F*_2, 58.526_ = 2.312, *P* = 0.108). Sex ratio showed a weak positive association with proximity to conspecifics (*F*_1, 10.802_ = 3.764, *P* = 0.079), meaning packs with a higher male sex ratio stayed closer together. There was also a weak positive effect of number of playback day, indicating an increase in proximity to conspecifics across sessions (*F*_1, 58.387_ = 3.454, *P* = 0.068). The full-null model comparison for gazing at group members was not significant (χ^2^ = 4.487, *P* = 0.344); therefore, individual predictors were not interpreted further.

## DISCUSSION

Chorus howling plays an important role in wolves’ social communication, shaping both intra- and inter-pack interactions. The results of this study show that zoo-housed wolves respond differently to synchronous and asynchronous chorus howls from unfamiliar packs. Following asynchronous howls, wolves were more active and showed investigatory responses directed towards the sound-source, including closer proximity, more frequent approaches, and longer gazing than following synchronous howls. Sex ratio also played a central role in how wolf packs responded to the unfamiliar stimuli.

Interestingly, behavioural responses to the dove control call were not consistently lower or clearly distinct from the responses to the howl conditions. Its intermediate response between synchronous and asynchronous playbacks in behaviours such as proximity to the sound-source or time spent moving could be interpreted as wolves’ behavioural reaction to a neutral stimulus, with more or less pronounced reactions to the howling playbacks. However, the high likelihood to approach the dove call as well as closer approach distance compared to the synchronized howl condition may suggest that the dove was not entirely perceived as a ‘neutral’ sound from the environment. This pattern may indicate that the dove call elicited some interest, whereby the wolves briefly inspected the sound-source without this leading to the same level of orientation towards the sound source as during the asynchronous playback. One possible explanation is that the presentation of the dove control call lacked ecological validity as it was broadcasted at an unnaturally high amplitude and from the ground rather than from elevated perches where doves typically call (Romagosa & Mlodinow, 2020). While this may have made the stimulus unusually salient or aversive to the wolves, we deemed the choices necessary to provide a direct control for the wolf howl playbacks. In line with that argument, it is also likely that aspects of the loudspeaker itself, such as the artificial quality of the sound transmission, may have contributed to the wolves’ responses (Deecke, 2006; Fischer et al., 2013), which is why we included the control in the first place. However, the fact that significant differences still emerged between the howl playback conditions despite a possible effect of the loudspeaker underlines that wolves indeed discriminate between chorus howls based on their synchrony.

Indeed, the response to synchronous howls partly supports the hypothesis that temporal vocal coordination may be perceived as more formidable in wolves (Bortolini et al., 2025; Fessler & Holbrook, 2016). Wolves remained farther from the sound-source, approached it less closely, less often, and reduced their activity following the synchronous compared with the asynchronous howls. This pattern may indicate a more cautious response to synchronous howls, potentially reflecting perceived threat or uncertainty. Highly coordinated acoustic signals can be perceived as more cohesive and impressive by receivers, and synchrony may therefore function as a cue of group strength (Bortolini et al., 2025; Fessler & Holbrook, 2016; Hall & Magrath, 2007). In addition, more synchronous or harmonious signals were shown to be harder to localise (Bregman, 1990; Harrington & Asa, 2003), which we hypothesised may have further increased uncertainty for the receiving pack. Indeed, wolves gazed and oriented directly towards the sound-source less often in the synchronous than the asynchronous condition, underlining that lack of certainty where exactly the sound came from as a potential explanation. However, we have to point out that since the responses to the dove call were more similar to the synchronous playback than to the asynchronous playback, the pattern observed in the synchronous condition could also be interpreted as lower interest or lower motivation to engage with the stimulus. Therefore, the interpretation that synchronous howls elicited a more cautious response due to perceived threat or uncertainty should be treated with caution.

In contrast, the responses to asynchronous howls did not support the prediction that ambiguity about the caller number would lead to stronger avoidance and increased intra-pack cohesion. Instead, wolves spent more time closer to the sound-source, approached it more often, moved more, oriented towards it more and gazed towards the source for longer following asynchronous howls compared to synchronous howls and mostly also compared to the dove call. This suggests that asynchronous howls led to increased arousal and sound-orientation, with a behaviour pattern that resembles an investigative rather than an avoidance response. In addition to the reduced certainty of where the sound came from in the synchronous condition, uncertainty about the caller number – as suggested by the Beau Geste effect (Harrington, 1989; Krebs, 1977) – may have thus likewise contributed to the increased orientation towards the sound source in the asynchronous condition. While we had expected avoidance in line with wild wolf behaviour towards threatening stimuli (Lazzaroni et al., 2026), in captive packs in particular, approach and investigative behaviour is often observed in response to threat-like stimuli such as passing dogs (pers. comm., Berweiler et al. in progress) or the scent marks of unfamiliar conspecifics (Studer et al., 2026; Wirobski et al., 2023). This could suggest that the asynchronous howls were perceived as more threatening than the synchronous ones and warranted investigation. A similar within-species effect has been reported in a recent study showing that humans responded more strongly to unfamiliar, asynchronous football chants than to synchronous chants, perceiving them as more threatening (Newson et al., in review). Interestingly, the same study found no such effect when humans listened to the same synchronous and asynchronous wolf howls used in this study, suggesting that this perception may be specific to species-relevant cues. In how far these effects in wolves and humans may be due to ambiguity in the caller number, as suggested by the Beau Geste effect (Harrington, 1989; Krebs, 1977), or whether the easier localisation of asynchronous compared to synchronous howls (Bregman, 1990; Harrington & Asa, 2003) played a role at least in the wolf study, remains to be explored in detail.

In addition to the effect of howl synchronization, sex ratio in the group also influenced the behavioural response. Groups with more males gazed more towards the sound-source and spent more time oriented towards it than groups with more females. At the same time, they also tended to move less and stayed closer together. This pattern may indicate that males were more attentive to unfamiliar howls, which is consistent with previous findings, suggesting that males, particularly dominant and experienced males, often play an important role in territory defence and inter-pack encounters (Cubaynes et al., 2014; Harrington & Mech, 1979; Mech & Boitani, 2003). We could also observe an effect of group size. Contrary to our expectations, larger packs remained at a greater distance from the sound-source (Cubaynes et al., 2014; Harrington & Asa, 2003; Harrington & Mech, 1979). However, group size correlated with enclosure size (see ary Material S7), therefore we do not interpret this effect further.

Some aspects should be considered when interpreting these findings. For one, the study was conducted with captive animals and needs to be evaluated with their life experience in mind. In contrast to free-ranging wolves, which use howling as an important component of territorial defence and interactions with neighbouring packs (Harrington & Mech, 1979, 1983), captive wolves experience limited opportunities for natural territorial interactions. As has also been suggested in studies using real animals as a potential external threat, the absence of experience with territorial intrusions in captivity may alter behavioural responses to simulated intruders (Berweiler, in progress; Studer et al., 2026). Nevertheless, the fact that the packs showed a significant behavioural difference following synchronous and asynchronous howling playbacks despite not having any previous experience with unfamiliar pack howls and their possible consequences, suggests that more inherent mechanisms may be at play. While the current study chose a captive setting to allow for controlled playback and observational conditions, future studies in wild populations would be valuable to assess whether the response patterns observed in captive wolves also occur under natural, ecologically more valid territorial conditions.

Secondly, the use of concatenated howl bouts as playback stimuli represents a compromise between ecological realism and experimental standardisation. Because field recordings of animal vocalisations are often limited in duration and affected by environmental noise, playback stimuli commonly require editing to obtain acoustically comparable exemplars (Fischer et al., 2013). To generate playback tracks of ecologically realistic duration while maintaining controlled differences in temporal coordination, the original howl recordings were extended through bout concatenation and separated by short silent intervals. These silent intervals also reflect the natural temporal organisation of wolf chorus howling, which typically comprises multiple howl bouts interspersed with brief pauses rather than uninterrupted vocalization (Harrington & Mech, 1979). Importantly, while we cannot exclude its influence on the animals’ behaviour, identical editing procedures were applied to both synchronous and asynchronous treatments, ensuring that any artefacts associated with stimulus construction were balanced across conditions and are therefore unlikely to account for differences in behavioural responses.

Third, three weeks were chosen as timeline between playbacks to maximise time between playback days while fitting the logistics of repeated testing at different locations into the targeted time of year outside of breeding and puppy season. Nevertheless, the effect of playback day number on sound approach, body orientation towards the sound-source, and proximity to conspecifics suggests that habituation may have been at play. Wolves can quickly habituate to stimuli they initially perceived as threatening, which has been shown even in wild wolves, though the same stimuli were used repeatedly in that study (Lazzaroni et al., 2026). Counterbalancing playback orders across packs, ideally with even longer (i.e., more than 3 weeks) between-stimuli intervals, will thus remain a central control factor for future playback experiments with wolves.

Fourth, the playback stimuli consisted of vocalisations from only two wolves. While previous research suggests that chorus howls may make the number of callers difficult to assess acoustically (Harrington, 1989), more recent studies found that acoustic indices of chorus howls correlate with the number of vocalising wolves, even though estimates from natural choruses may be imprecise (Papin et al., 2019). Thus, one may argue that the uncertainty regarding the perceived number of callers may be lower with playbacks from only two wolves compared to chorus howls from bigger packs. However, as most groups in our study were relatively small (mean group size ± SD = 4.21 ± 1.84), it is not unlikely that uncertainty about whether two or more wolves were howling may have influenced how the playbacks were perceived. At the same time, similar response patterns were also apparent in larger groups, suggesting that uncertainty about the number of callers may not fully account for the observed differences between playback conditions. Whether similar response patterns would also occur when chorus howls are produced by larger packs therefore remains to be investigated.

## CONCLUSION

The present study suggests that wolves are sensitive to synchrony modulations in chorus howls from unfamiliar packs and adjust their behaviour in terms of caution, approach, and exploration. Howl synchrony did not simply increase or decrease response intensity as we had initially hypothesised but rather resulted in distinct behavioural response patterns. Synchronous howls were associated with more distance and less movement, whereas asynchronous howls elicited more approach behaviour reflecting an investigation-response in captive wolf packs. While this could be interpreted as support for the Beau Geste effect in wolves (Harrington, 1989), more nuanced studies and replications with wild wolf packs will be needed to distinguish the roles of caller pack size ambiguity and localizability.

## Acknowledgements

We thank the participating zoos for their cooperation and for granting permission to conduct this study: Cumberland Wildpark Grünau, Natur- und Tierpark Goldau, Tiererlebnis Buchenberg, Tiergarten Schönbrunn, Tierpark Bern, Tierpark Biel,Tierpark Lange Erlen Basel, Tierpark Stadt Haag, Tierwelt Herberstein, Wildnispark Zürich Langenberg, Wildpark Bruderhaus Winterthur, Zoo et piscine des Marécottes, Zoo La Garenne, and Zoo Salzburg Hellbrunn. We are especially grateful to the animal care staff and zoo personnel for their logistical support and valuable information about the study animals. We also thank Sarah Vlasitz for providing the acoustic stimuli used in this study.

The author(s) declare no competing interests.

## Author contribution

Conceptualization: MN, EW, FR, SC; Data curation: VK, SC, PAS; Formal analysis: VK, SC; Funding acquisition: FR, GW, VK; Investigation: VK, SC; Methodology: VK, SC, FR; Project administration: VK, SC; Resources (NA); Software (NA); Supervision: FR, SC; Validation (NA); Visualization: VK, PAS; Roles/Writing - original draft: VK, SC; and Writing - review & editing: MN, EW, FR, PAS, GW

## Data availability

All data and analysis code necessary to reproduce the results reported in this article will be made available in the Figshare repository under the following link: https://doi.org/10.6084/m9.figshare.33295311

